# Endogenous epitope tagging of eEF1A2 in mice reveals early embryonic expression of eEF1A2 and subcellular compartmentalisation of neuronal eEF1A1 and eEF1A2

**DOI:** 10.1101/2023.04.20.537636

**Authors:** Faith C.J. Davies, Grant F. Marshall, Danni Gadd, Catherine M. Abbott

**Affiliations:** Centre for Genomic & Experimental Medicine, MRC Institute of Genetics and Molecular Medicine, University of Edinburgh, Western General Hospital, Crewe Road, Edinburgh EH4 2XU, United Kingdom; Simons Initiative for the Developing Brain, University of Edinburgh, Edinburgh EH8 9XD, United Kingdom

**Keywords:** epitope tagging, eEF1A1, eEF1A2, development, axons, translation elongation, neurodevelopment

## Abstract

All vertebrate species express two independently-encoded forms of translation elongation factor eEF1A. In humans and mice eEF1A1 and eEF1A2 are 92% identical at the amino acid level, but the well conserved developmental switch between the two variants in specific tissues suggests the existence of important functional differences. Heterozygous mutations in eEF1A2 result in neurodevelopmental disorders in humans; the mechanism of pathogenicity is unclear, but one hypothesis is that there is a dominant negative effect on eEF1A1 during development. The high degree of similarity between the eEF1A proteins has complicated expression analysis in the past; here we describe a gene edited mouse line in which we have introduced a V5 tag in the gene encoding eEF1A2. Expression analysis using anti-V5 and anti-eEF1A1 antibodies demonstrates that, in contrast to the prevailing view that eEF1A2 is only expressed postnatally, it is expressed from as early as E11.5 in the developing neural tube. Two colour immunofluorescence also reveals coordinated switching between eEF1A1 and eEF1A2 in different regions of postnatal brain. Completely reciprocal expression of the two variants is seen in post-weaning mouse brain with eEF1A1 expressed in oligodendrocytes and astrocytes and eEF1A2 in neuronal soma. Although eEF1A1 is absent from neuronal cell bodies after development, it is widely expressed in axons. This expression does not appear to coincide with myelin sheaths originating from oligodendrocytes but rather results from localised translation within the axon, suggesting that both variants are transcribed in neurons but show completely distinct subcellular localisation at the protein level. These findings will form an underlying framework for understanding how missense mutations in eEF1A2 result in neurodevelopmental disorders.

## Introduction

Neurons depend on protein synthesis not only for survival, but to enable a rapid response to environmental signals. Since the majority of mRNAs have a relatively long half-life, rapid changes in gene expression are more efficiently accomplished by changes in rates of protein synthesis and degradation. Translational control not only enables cells to respond rapidly to signalling, but for subsets of mRNAs to be translated in specific subcellular compartments. This system is particularly crucial for neuronal function; here the translational machinery is compartmentalised in the soma, dendrites and axons, and at synapses, permitting these often very physically distant compartments to respond to distinct signals (Holt *et al*., 2019). Localised translation in response to synaptic activity is essential for synaptic plasticity and memory consolidation, and axonal translation is now widely accepted as an important mechanism for establishment and maintenance of neuronal networks (Lin *et al*., 2021).

Translation elongation factor eEF1A plays a pivotal role in protein synthesis where it is responsible for the delivery of aminoacylated tRNAs to the ribosome (Merrick, 1992). This is a GTP dependent step, facilitated by the GTP exchange complex eEF1B (Le Sourd *et al*., 2006). In all known vertebrate species eEF1A exists as two independently encoded variants with distinct expression patterns; these variants are called eEF1A1 and eEF1A2. There is often confusion between the two in the literature due to the high degree of homology they exhibit at the amino acid level (92% identity, 98% similarity in human (Soares *et al*., 2009)). Most commercially available antibodies and many nucleic acid probes recognise both variant forms, complicating our understanding of their expression. However, multiple studies have shown that the eEF1A variants are differentially expressed in a developmental- and tissue-specific manner. Whilst eEF1A1 is expressed ubiquitously during development, and thereafter in almost all cell types, it becomes downregulated postnatally in brain and muscle in all vertebrates so far studied. In mice and rats eEF1A1 is downregulated to undetectable levels in muscle soon after weaning, at about 3 weeks of age. During this postnatal period, eEF1A2 becomes upregulated until it peaks at about 3 weeks and the switch is complete. In both mouse and rat this process seems to be regulated at the RNA level, as both mRNA and protein expression levels change in parallel (Lee *et al*., 1992; Chambers *et al*., 1998; Khalyfa *et al*., 2001).

In neurons, in contrast, the postnatal switch from eEF1A1 to eEF1A2 has been less well characterised, largely because expression analysis in brain is complicated by the fact that glia express eEF1A1 at high levels, rendering whole tissue analysis difficult. However, Westerns of whole mouse brain, *in situ* hybridisation and immunohistochemistry have been used to study expression of both variants, reaching the conclusion that in brain and spinal cord of post-weaning animals, eEF1A1 is no longer expressed in neurons but is replaced with the closely related variant eEF1A2 (Khalyfa *et al*., 2001; Pan *et al*., 2004a). Studies of early development have been limited to showing that eEF1A2 is not detectable by Western blotting in brain lysates at embryonic day 18 (Khalyfa *et al*., 2003), but can be detected in brain by immunofluorescence at postnatal day 1 (P1;(Pan *et al*., 2004b)). In contrast, eEF1A1 mRNA and protein have been shown to be the major form in prenatal and early postnatal neurons (Chambers *et al*., 1998; Khalyfa *et al*., 2001; Pan *et al*., 2004b), and throughout life in glia and white matter (Lee *et al*., 1995) (Newbery *et al*., 2007). Khalyfa et al reported a lack of expression of eEF1A1 in dendrites of postnatal animals (Khalyfa *et al*., 2003).

In spite of this earlier work, it is still unknown whether all neurons in adult animals, in every region of the brain, express only eEF1A2. Furthermore, numerous proteomic and transcriptomic analyses in both humans and rodents have demonstrated the presence of both eEF1A1 and eEF1A2 in synaptosomes, repeatedly identifying eEF1A1 as a synaptic mRNA or protein in both developing and fully mature organisms (Bayes *et al*., 2011; Bayes *et al*., 2012; Cajigas *et al*., 2012). The obvious caveat here is that eEF1A1 is strongly expressed in glia, which can contaminate postsynaptic density or neuropil preps, but this has largely been corrected for in the analyses performed (Cajigas *et al*., 2012). In addition, many more recent studies have reported expression of eEF1A1 in dendrites of adult animals, including Perez et al who showed dendritic enrichment of eEF1A1 mRNA in GABAergic interneurons (Perez *et al*., 2021).

The functional consequences of eEF1A variant switching during development are still unknown, though Mendoza et al (Mendoza *et al*., 2021) showed that phosphorylation of specific sites on eEF1A2 (that are not shared with eEF1A1) modulates structural plasticity in dendritic spines, coordinating protein synthesis with actin dynamics. One of these sites, Ser358, had previously been shown by Gandin et al to regulate stress-induced degradation of newly synthesised polypeptides, implying an additional role for eEF1A2 specifically in quality control of translation (Gandin *et al*., 2013). eEF1A1 is assumed to be an essential protein from early in development (since it has been shown to be essential in lower organisms like yeast (Cottrelle *et al*., 1985)), but mice with complete absence of eEF1A2 survive until ∼4 weeks postnatal, dying of motor neuron degeneration (Chambers *et al*., 1998; Newbery *et al*., 2005).

No humans with null mutations in eEF1A2 have been reported, but a wide range of *de novo* heterozygous missense mutations in the *EEF1A2* gene have been found to result in often severe neurodevelopmental disorders including early onset epilepsy and intellectual disability (Nakajima *et al*., 2015; Lam *et al*., 2016; Carvill *et al*., 2020). Affected children frequently also exhibit autistic behaviours and sleep disorders, and many have movement disorders or undergo a degenerative course (Carvill *et al*., 2020). The only known cases with homozygous mutations in *EEF1A2* were seen in one family where three children died before the age of 5 with dilated cardiomyopathy and severe epileptic encephalopathy (Cao *et al*., 2017). Recapitulation of pathogenic mutations in the mouse suggests that at least some of these missense mutations represent a gain of function (Davies *et al*., 2017; Davies *et al*., 2020).

A detailed description of the patterns of expression of the two eEF1A variants during brain development is key for understanding and treating disorders resulting from mutations in eEF1A2, but very little work has been published using two colour immunofluorescence for eEF1A1 and eEF1A2 simultaneously, largely due to issues with crossreactivity and the need to demonstrate antibody specificity. In order to address this gap in knowledge we introduced an epitope tag at the C terminus of the endogenous eEF1A2 protein in mice by gene editing, to allow us to detect eEF1A2 specifically when in combination with eEF1A1. We show that eEF1A2 is expressed from as early as E11.5 in some cell types, demonstrate the switch from eEF1A1 to eEF1A2 in different areas of the brain leading to non-overlapping expression patterns. Further, we show that in neurons eEF1A1 expression is not switched off throughout the cell, but rather switches from the soma to axons once development is complete.

## Materials and methods

### Transgenic mouse generation

Mice were housed in the Biomedical Research Facility (BRF) at the University of Edinburgh. All mice were maintained in accordance with Home Office regulations and all protocols had been approved by the local ethics committee of the University of Edinburgh. All methods were performed in accordance with the relevant guidelines and regulations. Embryo transfer was carried out with short term recovery anaesthesia and analgesia where needed post-operatively. Mutant mice were closely observed for overall clinical condition and were euthanized where necessary to avoid suffering.

Transgenic mice were made using the EASI-CRISPR method first described by Quadros et al (Quadros *et al*., 2017). gRNA (sequence TTCTTTAGTTAGACCAAACTAGG) was designed to cut the genome in intron 7 of *Eef1a2* to minimise chances of unwanted indels around the cut site affecting gene function. To insert the V5 tag into the 3’ end of *Eef1a2* a 720bp repair template was designed. As well as adding the V5 tag, the template made a number of other silent mutations; this was necessary as much of the locus proved too GC-rich to synthesise. *Eef1a2* intron 7 was replaced with human EEF1A1 intron 7 in the template, and the *Eef1a2* 3’UTR was replaced with the woodchuck hepatitis virus post-transcriptional regulatory element (WPRE) and SV40 polyA. Homology arms were 67bp long (see supplementary data figure 1A).

The microinjection mix was prepared with components (all from IDT) at the following concentrations: crRNA/tracrRNA complex at 27ng/ul, ssDNA megamer at 10ng/ul, Alt-R s.p. HiFi Cas9 nuclease V3 (50ng/ul). Freshly prepared microinjection mix was introduced by pronuclear injection into oocytes derived from C57BL/6JCr *mice*.

Mice were housed in single sex, mixed genotype groups of between 2 and 5 animals. Both male and female mice were used for expression analysis. All observational phenotyping was carried out blind to genotype.

### Genotyping

Ear notches were taken at 14 days after birth, DNA extracted and used for PCR genotyping, initially by direct sequencing when identifying founders, then after establishment of colonies by analysis of band sizes of PCR products on a 1% agarose gel where a wild-type allele gave a band size of 176bp and a V5 knock-in 762 bp. PCR primers used were 5′–3′ TCTCTTTTGAAGAAGAACGCCT and GCTTAAGCTGTTCTGACCGT and PCR was carried out with annealing temperature of 62°C using Platinum Superfi polymerase (Thermofisher) for 34 cycles.

### Protein analysis

Whole brains and hind limb muscles were dissected, frozen on dry ice and stored at −70°C until use. They were homogenized in 10 μl/mg tissue 0.32 M sucrose containing protease inhibitor (cOmplete Mini, Roche) then centrifuged at 17000 g at 4°C for 30 min. The supernatant was removed and mixed with 4× NuPAGE LDS sample buffer and 10× NuPAGE reducing agent, then heated to 70°C for 10 min. Lysates were loaded onto NuPAGE 4–12% bis-tris gels and run at 200 V in NuPAGE MOPS SDS, before being transferred onto PVDF membrane (Amersham Hybond) in NuPAGE transfer buffer. Total protein was visualized using Revert solution (LI-COR). Blots were then blocked in Odyssey blocking buffer (LI-COR) for 1 h, incubated in either V5 anti-mouse (Invitrogen 46-0705) 1:2500 or 1:5000 in an antibody to eEF1A2 (custom made by Proteintech, equivalent to eEF1A2-1 in Newbery et al (Newbery *et al*., 2007)), then washed in TBS buffer containing 0.01% tween-20 for 3× 15 min, and incubated in IRdye 800CW anti-rabbit or anti-mouse (LI-COR) secondary antibody for 1 h. After another set of washes, images were taken using the Odyssey CLx (LI-COR) and analysed on Image Studio Lite Version 5.2.

### RNA analysis

Brain tissue was homogenised in Qiazol using a bead mill homogeniser (Precellys). RNA was isolated using the QIAGEN RNeasy mini kit and treated with QIAGEN RNase-Free DNase Set. RNA quality was assessed using a Bioanalyzer, and only RNA with a RIN above 7.8 was used for cDNA synthesis. cDNA was then synthesized using the Agilent Technologies AffinityScript Multi Temperature cDNA Synthesis Kit. Quantitative RT-PCR (qPCR) using primers for eEF1A2 exon 2–4 (5′ GCCACGATCAGCACTGCG and 5′ CAAGCGGACCATCGAGAAGT), eEF1A2 exon 3–4 (5′ ATATGATTACAGGCACATCCCAG and 5′ GTGCGTGTTCCCGGGTTT). GAPDH (5’ GGAAGGGCTCATGACCACA and 5’ CCGTTCAGCTCTGGGATGAC) and UBC (5’ AGCCCAGTGTTACCACCAAG and 5’ ACCCAAGAACAAGCACAAGG) were selected as suitable reference genes after analysis with a geNorm 6 gene mouse kit (PrimerDesign). Brilliant II SYBR Green QPCR master mix (Agilent) was used to make the reaction mix. cDNA was diluted 1:10 and 4 μl added to each reaction. Samples were run in triplicate in a LightCycler HT7900 (Roche) and quantity means were calculated using Ct values along with the slope and the Y-intercept from the standard curve for each gene. Statistics were performed using t-tests in GraphPad Prism V8.

### Timed matings

Proven males and 8-12 week old females were co-housed in the late afternoon. Females were checked for the presence of vaginal plug every subsequent morning, and weight of dam recorded once a plug was found. The morning the plug was found was taken to be day 0.5 post-fertilisation. Weight gain of >1.75g on day of collection indicated probable pregnancy.

### Immunofluorescence

Mice were humanely killed and tissues taken and rinsed in ice-cold PBS and fixed overnight at 4°C in at least 15 volumes of 4% paraformaldehyde. Tissues were stored in 50% ethanol until processing by dehydration and paraffin embedding, and were then cut into 10um sections and mounted on glass slides. Slides were deparaffinised with xylene, rehydrated and microwaved in citric acid pH6 for 20 minutes. They were then blocked for 1 hour in a 1:5 dilution in PBS of sheep or donkey serum, using the species in which the secondary antibody was raised. Primary antibodies were applied in 1% serum in PBS with 0.1% triton-X and incubated overnight at 4°C. After 3 PBS washes slides were treated with 1x TrueBlack or 2x TrueBlack Plus (Biotium) as per manufacturer’s instructions, then incubated with secondary antibody diluted 1:500-1:1000 in 1% animal serum in PBS for one hour. After another 3 PBS washes slides were coverslipped using Vectashield Vibrance + DAPI and left to cure overnight at room temperature.

Epifluorescent images were acquired using either A: a Zeiss Axio Scan Z1 slide scanning microscope (Carl Zeiss UK, Cambridge, UK) with Fluar or Plan Apochromat objective lenses (2.5x, 5x, 10x, 20x, 40x), a Zeiss Colibri 7 LED light source and a Zeiss Axiocam 506m monochrome CCD camera and Zeiss 90 HE, 92 HE, 96 HE, 38 HE and 43 HE fluorescent filter sets, with image capture performed using Zeiss Zen 3.5 Slidescan acquisition software. Or B: a Photometrics Prime BSI CMOS camera (Photometrics, Tuscon, AZ) fitted to a Zeiss AxioImager M2 fluorescence microscope with Plan-Apochromat objectives, using either 1. A Zeiss Colibri 7 LED light source, together with Zeiss filter sets 90 HE, 92 HE, 96 HE, 38 HE and 43 HE (Carl Zeiss UK, Cambridge, UK), with image capture performed in Zeiss Zen 3.5 software, or 2. A Mercury Halide fluorescent light source (Exfo Excite 120, Excelitas Technologies) and Chroma #89014ET three colour filter set (Chroma Technology Corp., Rockingham, VT), with single excitation and emission filters installed in motorised filter wheels (Prior Scientific Instruments, Cambridge, UK) and image capture performed using Micromanager (Version 1.4).

Confocal images were acquired using a Multimodal Imaging Platform Dragonfly (Andor technologies, Belfast UK) equipped with 405, 445, 488, 514, 561, 640 and 680nm lasers built on a Nikon Eclipse Ti-E inverted microscope body with Perfect focus system (Nikon Instruments, Japan). Data were collected in Spinning Disk 40μm pinhole mode with a 100x objective on the iXon 888 EMCCD / Zyla 4.2 sCMOS camera using a Bin of 1×1 and frame averaging of 1 using Andor Fusion acquisition software.

### Antibodies

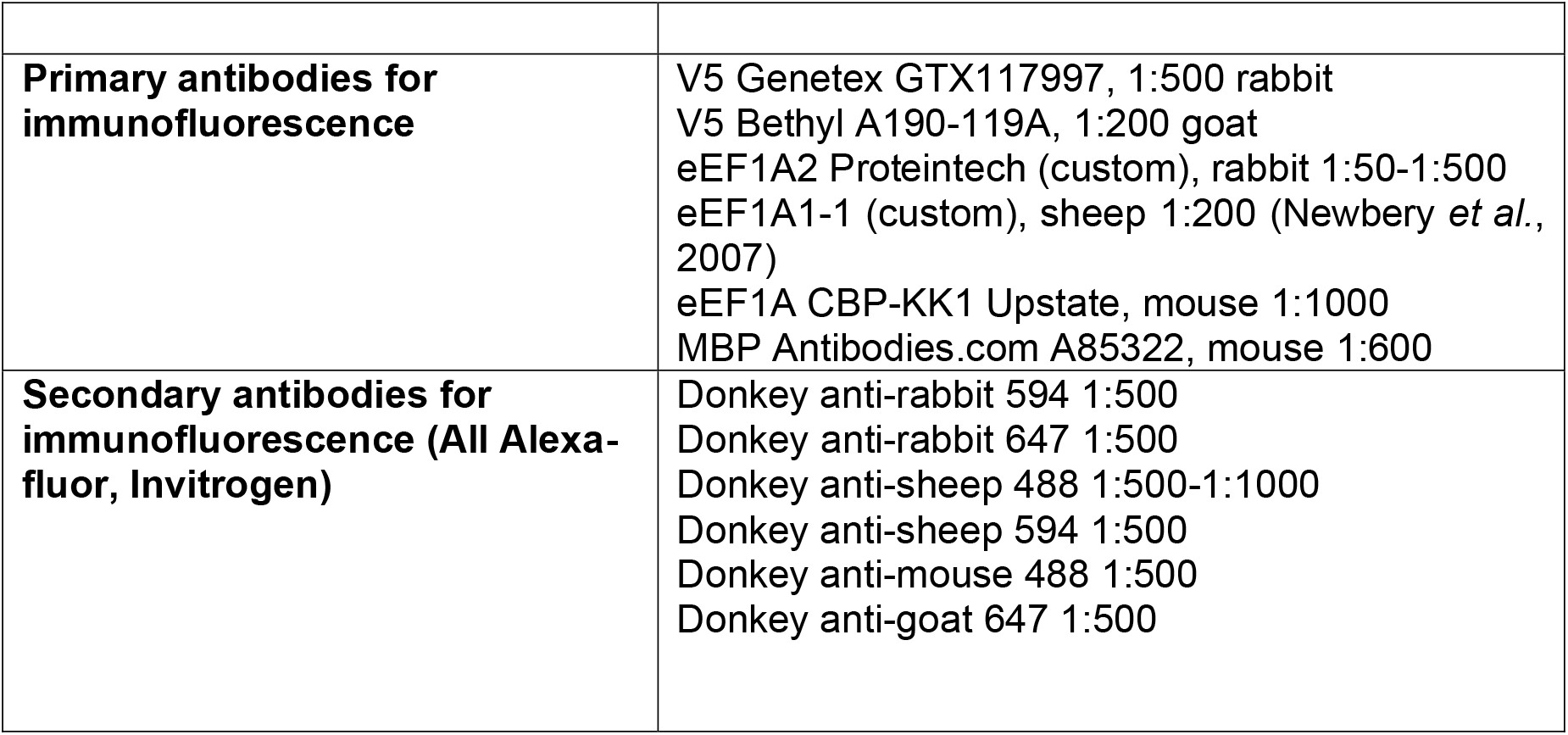

|  |  |
| --- | --- |
| **Primary antibodies for immunofluorescence** | V5 Genetex GTX117997, 1:500 rabbit<br>V5 Bethyl A190-119A, 1:200 goat<br>eEF1A2 Proteintech (custom), rabbit 1:50-1:500<br>eEF1A1-1 (custom), sheep 1:200 (Newbery *et al.*, 2007)<br>eEF1A CBP-KK1 Upstate, mouse 1:1000<br>MBP Antibodies.com A85322, mouse 1:600 |
| **Secondary antibodies for immunofluorescence (All Alexa-fluor, Invitrogen)** | Donkey anti-rabbit 594 1:500<br>Donkey anti-rabbit 647 1:500<br>Donkey anti-sheep 488 1:500-1:1000<br>Donkey anti-sheep 594 1:500<br>Donkey anti-mouse 488 1:500<br>Donkey anti-goat 647 1:500 |

## Results

### Engineering an epitope tag into the mouse Eef1a2 gene

Differential expression analysis of eEF1A1 and eEF1A2 is non-trivial using standard antibodies because of the high chance of cross-reactivity resulting from the 92% identity between the two proteins. We therefore opted to introduce a V5 epitope tag at the 3’ end of the endogenous *Eef1a2* mouse gene using CRISPR/Cas9 gene editing, with the aim of producing a line of mice expressing eEF1A2 protein with a C-terminal V5 tag that could be specifically detected in immunofluorescence. We designed a single-stranded oligonucleotide repair template of 720 nucleotides that included the whole coding portion of exon 8 of the *Eef1a2* gene with a V5 tag in-frame. This was followed by the WPRE element (which is widely used in gene therapy and here used as a replacement for the 3’UTR of eEF1A2, which could not be synthesised), all flanked by homology arms; see Figure 1A and supplementary information. It also proved impossible to synthesise intron 7 so this was replaced by the less GC-rich intron 7 from eEF1A1. The repair template was introduced into fertilised oocytes together with pre-assembled crRNA + tracrRNA + Cas9 ribonucleoprotein (ctRNP) complexes using Easi-CRISPR (Quadros *et al*., 2017) and the resulting live-born mice genotyped by sequencing. Of 27 founders born, 16 had incorporated the repair template on one allele and 8 were homozygous for the V5 tag with only 8 mice showing no indication of a knock-in.

**Figure 1.**
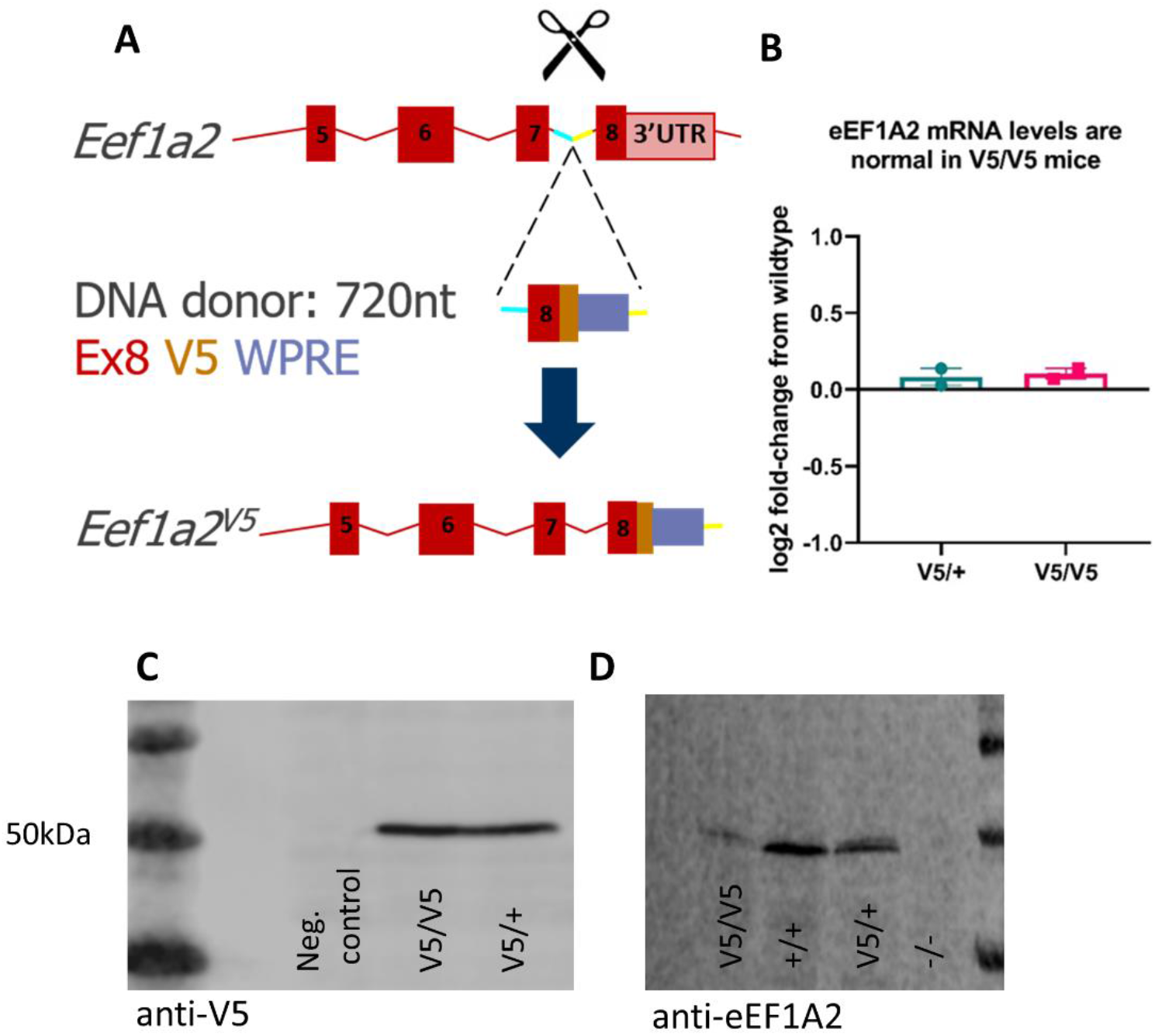
A) Schematic of the CRISPR design used to create the eEF1A2-V5 mouse line, with the cut site indicated by scissors and the V5 and WPRE parts of the repair template in orange and blue respectively. Exons are shown as boxes, with solid boxes correspond to coding regions. B) qPCR results showing that mice both heterozygous and homozygous for the V5 insertion express similar levels of eEF1A2 mRNA to those seen in WT mice. C) Western blot showing a single band of the correct size detected in brain with an anti-V5 antibody in heterozygous and homozygous mice. D) Western blot showing that the V5 tagged eEF1A2 protein is detected by an anti-eEF1A2 antibody but with reduced expression in V5/V5 homozygous mice.

A line was generated from one of the mice heterozygous for the knock-in and expression analysis performed. Whilst the levels of tagged eEF1A2 mRNA were equivalent to those in wild-type (WT) littermates (Figure 1B) in both heterozygous and homozygous V5 mice, Western blotting showed clearly that V5 carrying mice were producing a single band of the correct size, whether detected with an anti-eEF1A2 or anti-V5 antibody, although levels did appear to be reduced (Figure 1C, D). Homozygous knock-in mice died by ∼4 weeks, at a similar time to eEF1A2-null mice (Chambers *et al*., 1998) consistent with the low levels of expression seen. It is unclear whether the tag or the replacement of the 3’UTR with the WPRE element caused the reduction in expression, but the normal mRNA levels suggest that the tag might interfere with protein stability. Nevertheless, the heterozygotes were phenotypically normal in terms of weight and survival and expression of the tagged protein was sufficiently high for the resulting line to be used for assessing expression of eEF1A2 in tissues throughout development. These mice are referred to as V5/+ (heterozygotes) and V5/V5 (homozygotes) throughout, for ease of reading.

### Expression of the tagged eEF1A2 protein reflects that of endogenous eEF1A2

Previous work using immunofluorescence imaging studies have showed eEF1A2 and eEF1A1 to be expressed reciprocally in different cell types of the brain. Broadly speaking, eEF1A2 is found in neuronal cell bodies, and eEF1A1 in glial cells and white matter tracts (Pan *et al*., 2004a; Newbery *et al*., 2007). In contrast, RNA-seq studies have shown both eEF1A1 and eEF1A2 transcripts in neuronal cells, indicating that the expression patterns of eEF1A isoforms in brain may be more nuanced and complex than first thought (Bayes *et al*., 2011; Bayes *et al*., 2012; Cajigas *et al*., 2012).

We initially carried out immunofluorescence on sections of the cerebellum as Purkinje cells (PC) are known to express high levels of eEF1A2 (Newbery *et al*., 2007). We examined expression of V5-tagged eEF1A2 in the cerebellum of post-weaning mice. Staining with an anti-V5 antibody revealed strong, specific expression in PC in heterozygous (V5/+) mice, and equally specific expression, albeit at lower levels, in V5/V5 animals (Figure 2A). Co-immunofluorescence with anti-eEF1A2 and anti-V5 antibodies showed completely overlapping expression in PC of a V5/V5 mouse. There were some minor areas of non-specific background staining with the anti-V5 antibody in non-neuronal areas but these were also seen in the WT mouse not expressing V5; Figure 2B. Staining of neuronal cell bodies also revealed overlapping expression of the tagged and total eEF1A2 protein at the subcellular level (Figure 2C).

**Figure 2.**
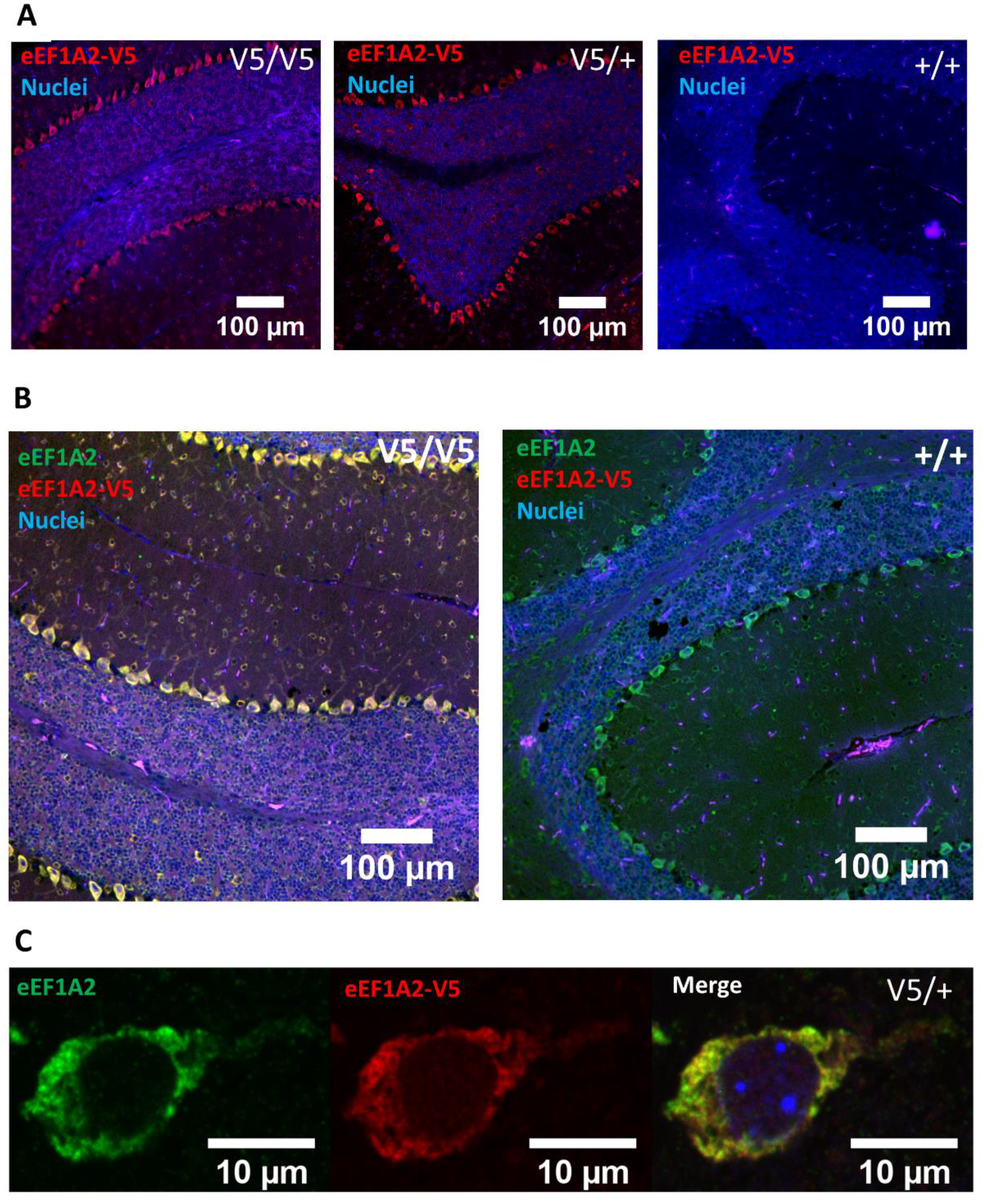
A) Immunofluorescence images of the cerebellum of homozygous (V5/V5), heterozygous (V5/+) and WT mice stained with anti-V5 antibody showing reduced expression in V5/V5 mice and no detectable expression in the WT mouse. B) as above but co-immunofluoresence using anti-V5 and anti-eEF1A2 antibodies to show completely overlapping expression of eEF1A2 and V5 staining in the V5/V5 mouse but eEF1A2 only in the WT control (scalebars on A and B are 100μm). C) Confocal micrographs showing overlapping expression of eEF1A2 and V5 staining within the cell body of a cortical neuron.

### eEF1A2 is expressed as early as E11.5

We then examined expression of eEF1A2-V5 in V5/+ mice between the ages of E11.5 and P25. Timed matings in the V5 tagged line were used to collect embryos for sectioning at different stages. Although previous reports in the literature have suggested that eEF1A2 is not expressed until after birth, Figure 3A shows that as early as E11.5, V5-tagged eEF1A2 is detectable at low levels in the basal plate of the embryonic spinal cord. By E13.5 (Figure 3B) there is more widespread expression throughout the spinal cord and brain, with strong expression seen in heart, dorsal root ganglia, trigeminal ganglia and olfactory bulb.

**Figure 3.**
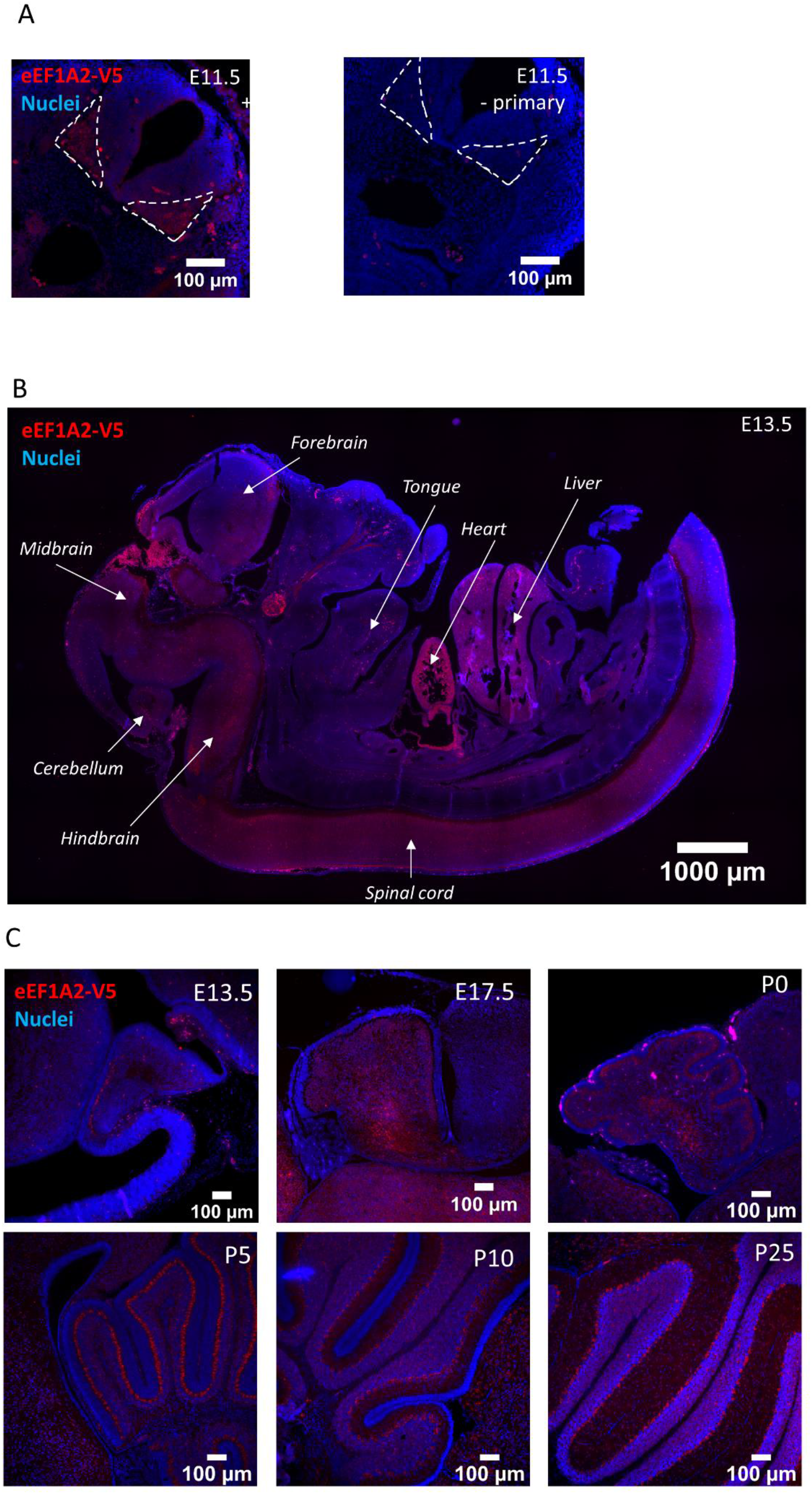
A) Immunofluorescence of a transverse section of an E11.5 embryo stained with anti-V5 antibody showing expression of V5-tagged eEF1A2 in the mantle layer of the basal plate of the developing spinal cord (white dashed lines). B) More widespread expression of V5-tagged eEF1A2 by E13.5. C) Sections of cerebellum at different stages, all stained with anti-V5 antibody, showing developing expression of eEF1A2 in Purkinje cells. Nuclei stained blue with DAPI in all images.

Analysis of V5 staining in the cerebellum at embryonic and postnatal stages shows eEF1A2 expression developing in Purkinje cells (PC). We show that eEF1A2 is found in cerebellar cell types as early as E13.5, around the time when PC are first differentiated in mouse (E11-E13) (Sotelo, 2004) and track the migration of PC through the developing cerebellum (Figure 3C).

### Non-overlapping expression of eEF1A1 and eEF1A2 throughout the brain by P25

We next examined expression of eEF1A2 throughout the brain at P25. Figure 4A shows mouse brain at P25 with eEF1A2 detected by anti-V5. Strong expression of eEF1A2-V5 can clearly be seen in neurons of multiple brain regions by this age. eEF1A2 is clearly expressed throughout the brain, with staining throughout the cortex, and is particularly strongly expressed in hippocampus and cerebellum. However, whilst expression of eEF1A2 is generally widespread, expression is low in the striatum. In order to establish whether this low staining level could be an artefact resulting from the tagging of eEF1A2 we carried out immunofluorescence using a commercial antibody which detects both eEF1A1 and eEF1A2 (Figure 4B). This shows clearly that in fact expression of either form of eEF1A is markedly lower in the striatum than in the surrounding areas of the brain, consistent with the findings of Ayola et al who found reduced rates of protein synthesis in the striatum (Avola *et al*., 1988).

**Figure 4.**
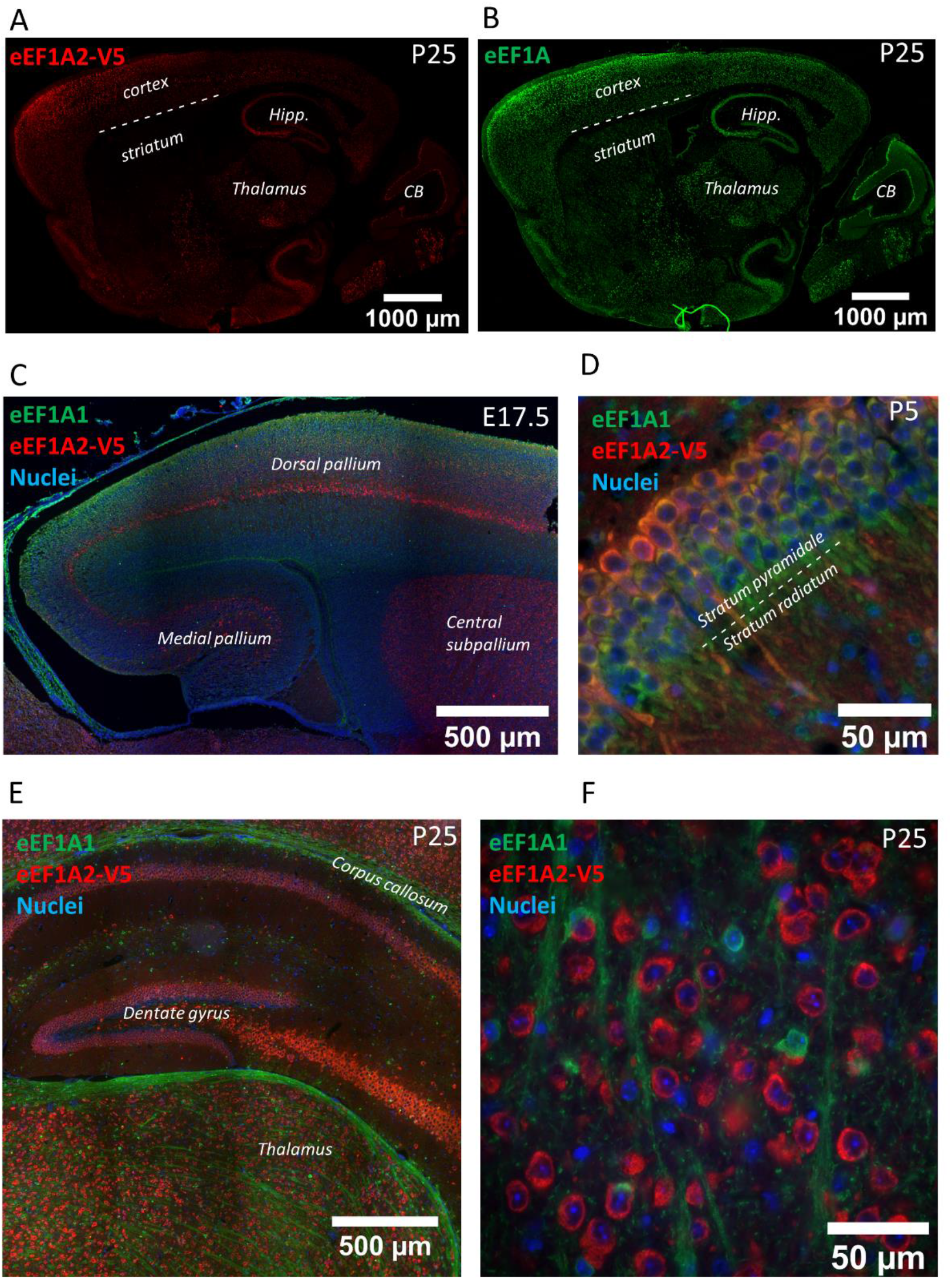
A) Composite image of whole brain of a V5/+ mouse at P25 stained with anti-V5 and B) a pan-eEF1A antibody that detects both eEF1A1 and eEF1A2, to demonstrate reduced expression of eEF1A in the striatum. C) Co-immunofluoresence of eEF1A1 and V5-tagged eEF1A2 (stained with anti-V5 antibody) shows predominant expression of eEF1A1 in the pallium at E17.5 with emerging D) Co-immunofluoresence of eEF1A1 and V5-tagged eEF1A2 (stained with anti-V5 antibody) shows a gradient of switching from eEF1A1 to eEF1A2 in the hippocampal pyramidal layer at P5.C) E) Co-immunofluorescence of eEF1A2-V5 and eEF1A1 in the hippocampus of a P25 mouse expressing the V5 tagged form of eEF1A2, showing non-overlapping expression patterns of the two variant forms of eEF1A. F) Co-immunofluorescence of eEF1A1 and V5-tagged eEF1A2 in thalamus at P25 showing clear staining of eEF1A1 in axons.

Co-immunofluorescence of V5-tagged eEF1A2 and endogenous eEF1A1 reveals changing patterns of the two variant forms of eEF1A in neurons during development. Figure 4C shows a section of hippocampus at E17.5 with predominant staining of eEF1A1 throughout (particularly in presumed neuronal progenitor cells) but with a focused band of strong staining of eEF1A2-V5 in the differentiating CA neurons. In figure 4D a section of hippocampus from mice aged P5 can be seen: here eEF1A1 is evenly expressed throughout pyramidal cells of CA3, whilst eEF1A2 shows a gradient of expression across deep and superficial layers, with stronger expression in the deep (more differentiated) pyramidal cells. By 25 days postnatal (figure 4E), analysis of expression of eEF1A1 and eEF1A2-V5 in hippocampus shows a dense band of expression of eEF1A2 in dorsal granule and pyramidal cells, and in dorsal CA1, CA2 and CA3 layers. By this stage of development there is no apparent overlapping expression of the two eEF1A variant proteins and eEF1A1 expression appears to be confined to axons under the cortex. A similarly mutually exclusive pattern of expression is seen in other brain areas (e.g. thalamus, figure 4F), with clear evidence of eEF1A1, but not eEF1A2, in axons.

### eEF1A1 protein expression is confined to axons in mature neurons

It is clear from the literature and the results above (figures 4E and F) that whilst eEF1A2 is highly expressed in neuronal soma, eEF1A1 is not expressed in mature neuronal cell bodies but is highly expressed in glial cells and white matter, including axon tracts. Since eEF1A1 is highly expressed in oligodendrocytes, it would be reasonable to assume that the staining seen in axons represents staining of the myelin sheath originating from oligodendrocytes. We set out to test this directly by performing co-immunofluorescence of eEF1A1 and myelin basic protein (MBP), a key component of the myelin sheath. Multiple sections through axon bundles in the cortex and striatum show clearly that anti-eEF1A1 and anti-MBP antibodies both stain axons (figure 5 A-D).

**Figure 5.**
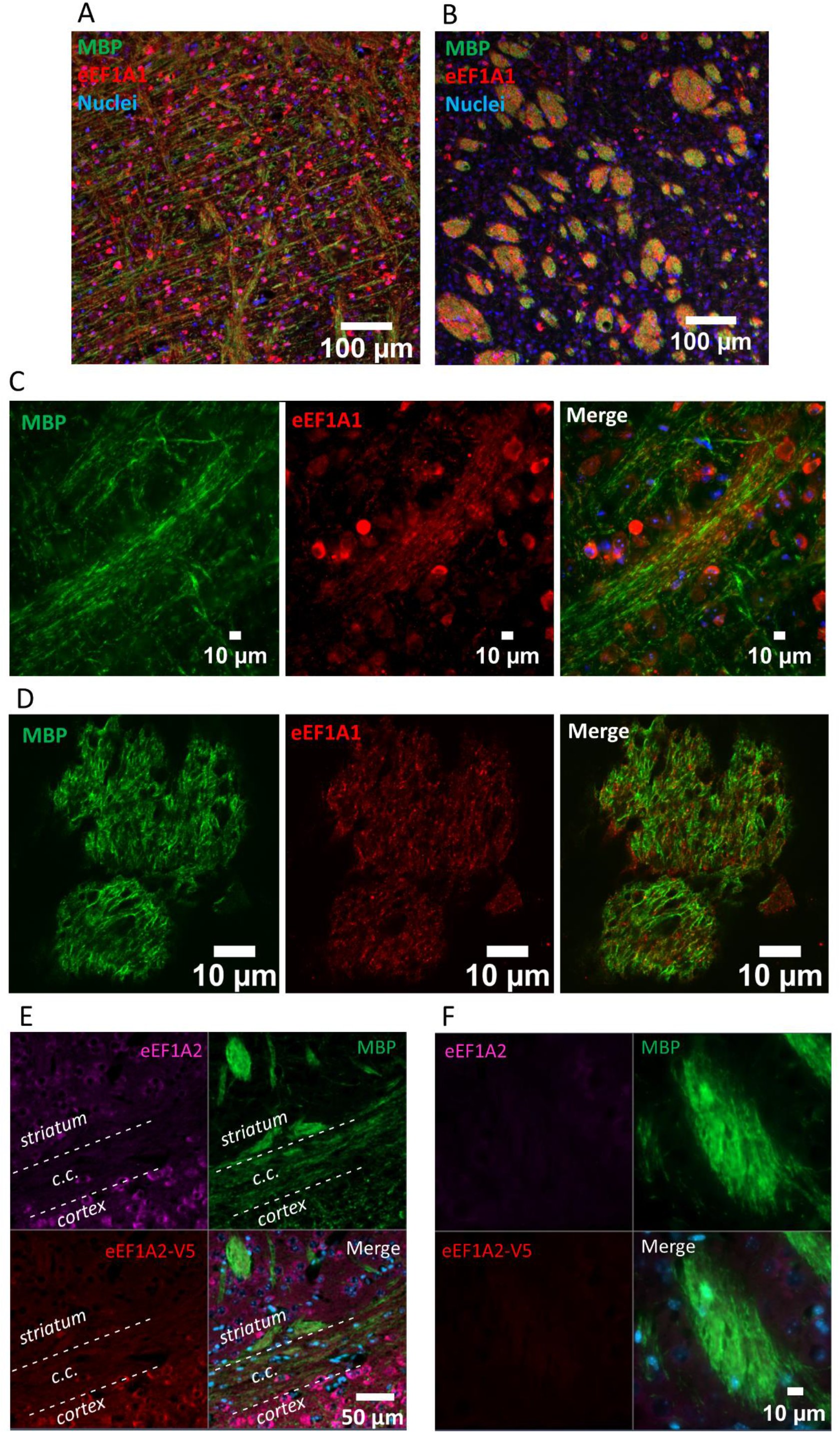
A, B) Co-immunofluorescence of eEF1A1 and MBP shows largely non-overlapping staining in the cortex (A) but overlapping staining in the straitum (B). C) Co-immunofluorescence of eEF1A1 and MBP shows co-localisation of eEF1A1 in cortical axons marked by MBP. D) Confocal micrographs showing overlapping and non-overlapping staining of eEF1A1 and MBP within subcortical axon bundles. E, F) Co-immunofluoresence of total eEF1A2 and V5-tagged eEF1A2 together with MPB shows the absence of both tagged and endogenous eEF1A2 in transverse and lateral sections through axons. The striatum, cortex and corpus callosum (c.c.) are labelled in panel E.

Due to poor structural preservation of axons in formalin-fixed paraffin embedded sections, we were unable to fully resolve whether eEF1A1 was within axons. However, it was clear that there were areas of both overlapping and non-overlapping staining of eEF1A1 and MBP in transverse sections through subcortical axon bundles (figure 5 D), suggesting that some eEF1A1 signal originated from within axons and not from non-neuronal cells.

Although we saw no overlapping expression between eEF1A1 and V5-tagged eEF1A2 in mature neurons, we wanted to exclude the possibility that the presence of the epitope tag had prevented eEF1A2 from being trafficked into axons. We carried out further immunofluorescence on sections from V5/+ mice (which therefore express both tagged and untagged eEF1A2) using three antibodies: anti-V5, anti-eEF1A2, and anti-MBP to stain the myelin sheath. Figures 5E and F show the absence of both tagged and untagged eEF1A2 within axons.

## Discussion

Epitope tagging of endogenous eEF1A2 has enabled us for the first time reliably to track the expression patterns of both eEF1A1 and eEF1A2 through development. This approach allowed us to circumvent issues of cross-reactivity of antibodies raised against the two closely related proteins, and the results form a valuable framework to underpin further studies investigating the functional consequences of the developmental switching from eEF1A1 to eEF1A2 in specific cell types and/or subcellular compartments. The EASI-CRISPR technique used to introduce the V5 tag worked extremely efficiently and could be applied to other proteins where antibody specificity cannot be achieved. We confirmed that staining with an anti-V5 antibody coincided perfectly with that seen using an antibody raised against eEF1A2, and that, crucially, the anti-V5 antibody was able to be used in co-immunofluorescence experiments with anti-eEF1A1.

We detected expression of eEF1A2 at E11.5, the earliest stage at which we looked. Expression was seen in heart and the central nervous system, increasing throughout development, and at later stages waves of switching between eEF1A1 and eEF1A2 could be observed in specific areas of brain, most notably in the cerebellum and hippocampus. In more mature brain, from postnatal day 25, a clear picture of non-overlapping expression of eEF1A1 and eEF1A2 emerges.

When stitched images of the whole brain stained with an antibody that detects both eEF1A1 and eEF1A2 are examined, it is clear that striatum shows very low levels of expression of both variants in relation to other areas of the brain (Figure 4A). The striatum is largely made up of medium spiny neurons which are GABAergic and have unmyelinated axons. A study of protein synthesis rates in 21 different areas of mouse brain found that the lowest levels were seen in corpus callosum, closely followed by striatum (Qin *et al*., 2011). This correlates with the low levels of eEF1A expression, though the cause and result of this variation in expression across different regions of the brain has yet to be established.

Some regions of strong expression of eEF1A2 may explain the phenotypes seen in individuals with heterozygous missense mutations in *EEF1A2*. Clearly the high levels of expression in cortex and hippocampus correlate well with the known epilepsy and intellectual disability phenotypes, but we also see very strong expression in Purkinje cells in the cerebellum and dorsal root ganglia. A number of children with mutations in *EEF1A2* who are able to walk have been found to have gait abnormalities such as ataxia; loss of Purkinje cells is well established to result in ataxia in both animal models and humans and has been often been shown to be associated with defects in tRNA metabolism (Lee *et al*., 2006; Kapur *et al*., 2017). The expression of eEF1A2 in heart from early in development may additionally explain the fatal dilated cardiomyopathy seen in children homozygous for the P333L missense mutation in *EEF1A2* (Cao *et al*., 2017), particularly in view of the recent report implicating eEF1A2 in cardiomyocyte differentiation (Lyu *et al*., 2022).

In addition to evidence for the mutually exclusive expression of eEF1A1 and eEF1A2 in the cell bodies of different cell types in the brain, we now show that eEF1A1 expression is not, as previously thought, shut down completely in mature neurons. Instead, it relocates from the neuronal soma, where it is highly expressed during embryonic and early postnatal development, to axons. This switching mechanism is in contrast to that seen in muscle where eEF1A1 is shut down transcriptionally around weaning in mice (Chambers *et al*., 1998; Khalyfa *et al*., 2003). Rather, transcription is likely sustained at similar levels throughout development, but either the eEF1A1 mRNA or protein changes localisation. Although proteins can be directly targeted to specific subcellular locations by virtue of specific motifs, protein localisation in neurons is frequently controlled by targeting of the cognate mRNA, which is then translated locally, and this seems the most likely mechanism here. Indeed, a number of studies of both the mRNA content and translatome of axons have identified eEF1A1 but not eEF1A2 in axons of mice and rats (Taylor *et al*., 2009; Gumy *et al*., 2011; Shigeoka *et al*., 2016; Nijssen *et al*., 2018) and in dendrites, albeit in cultured cells (Wefers *et al*., 2022); other studies using TRAP and Riboseq methodology have shown enrichment of eEF1A1 in axons relative to cell bodies (Ostroff *et al*., 2019; Glock *et al*., 2021; Jung *et al*., 2023). It seems likely that there are one or more RNA binding proteins controlling the developmental switch resulting in the transport and subsequent translation of eEF1A1 in axons; the UTRs of eEF1A1 and eEF1A2, unlike the coding regions, are largely dissimilar (41% identity in 3’UTRs compared with 80% in the coding regions).

A key question is why this switch between near-identical proteins exists, and what are the functional consequences of the change in subcellular localisation of eEF1A1 during neuronal maturation. Biochemically, the two variants have indistinguishable activity in an *in vitro* translation system, although with different affinities for GTP (eEF1A2 shows a greater affinity for GDP and eEF1A1 a greater affinity for GTP (Kahns *et al*., 1998)). The differences between eEF1A1 and eEF1A2 are highly conserved throughout vertebrate evolution, suggesting strong constraints on the amino acids that differ between variants, and a functional role for these differences. Nevertheless, the extent to which there is functional equivalence of the two isoforms, and the degree to which their activities differ, is still largely a mystery. However, some progress has been made by focusing on specific sites of variation between eEF1A1 and eEF1A2. At a number of locations, particularly at the C terminal end of eEF1A, the amino acid differences between the two variant forms are alanine (eEF1A1) to serine (eEF1A2) substitutions, raising the possibility of eEF1A2-specific phosphorylation (Soares *et al*., 2009). Indeed, Gandin et al (Gandin *et al*., 2013) demonstrated a role for serines 205 and 358 in regulating the degradation of newly synthesised proteins under conditions of stress, during which these serines are phosphorylated by the kinase JNK. Romaus Sanjurjo et al found that overexpression of eEF1A2, but not eEF1A1, in corticospinal tract neurons increased protein synthesis and actin rearrangements, leading to enhanced sprouting after experimentally induced neuronal injury (Romaus-Sanjurjo *et al*., 2022). Further, Mendoza et al (Mendoza *et al*., 2021) used phosphomimetic and nonphosphorylatable mutant forms of eEF1A2 transfected into hippocampal neurons to show differential effects of the mutant forms on actin binding and translation activity, suggesting a key role for eEF1A2 in coordination of translation and actin dynamics in dendritic spines in response to metabotropic glutamate receptor signalling. Since this activity depends on the serines that are present in eEF1A2 and not eEF1A1, it is reasonable to assume that the two variant forms of eEF1A will have distinct roles in the different subcellular compartments, and manipulation of the two, targeting them to different regions, will be necessary to understand their distinct, specific functions in axons, dendrites, and at synapses (where there is evidence for both variant forms, though not necessarily within the same synapse).

Our results have implications for increased understanding of the ways in which missense mutations in eEF1A2 can result in often severe neurodevelopmental disorders. There is evidence from both the unusual mutational profile of the gene, and from mouse models in which the effects of missense mutations are compared with null mutations on the same background (Davies *et al*., 2020), that the pathogenic variants result in some form of gain of function. The co-expression of eEF1A1 and eEF1A2 during development, notably in neuronal soma, suggests the possibility that missense mutant forms of eEF1A2 could exert a dominant negative effect on eEF1A1, particularly since there is evidence that eEF1A1 and eEF1A2 can heterodimerise (Sanges *et al*., 2012; Crepin *et al*., 2014). A mutation that affects the balance between monomers and homo- or heterodimers could in turn affect the balance in the cell between translation elongation (carried out by monomers) and cytoskeletal remodelling (performed by eEF1A dimers) (Bunai *et al*., 2006)). Such a disturbance could ultimately affect spine maturation, remodelling, and/or pruning, leading to changes in circuit activity and ultimately to epilepsy and intellectual disability. Indeed, the clear compartmentalisation of eEF1A1 and eEF1A2 within the mature neuron might suggest the need to avoid heterodimerisation once development is complete. It will be interesting to establish how widespread this compartmentalisation is in species that have different variant forms of eEF1A. Ultimately, functional studies will be needed to examine the role of higher order complexes of eEF1A on neuronal function and dysfunction.

Finally, the strategy of endogenous tagging that allowed us to circumvent problems with antibody specificity could be applied to other proteins where similar issues exist and where detailed mapping of protein level expression is necessary to understand disease processes.

## Abbreviations

eEF1A1/eEF1A2: eukaryotic translation elongation factor 1A1/2
WPRE: woodchuck hepatitis virus post-transcriptional regulatory element
PC: Purkinje cells
CA1-3: cortical area 1-3

## Acknowledgements

We are grateful to Professor David Price for critical reading of the manuscript and advice on analysis. The authors gratefully acknowledge the BVS Central Transgenic Core, the staff of the Biomedical Research Facility and the staff of the IGC Advanced Imaging Resource of the University of Edinburgh for their expertise and assistance in this work. We would like to dedicate this paper to the memory of Emma Allan, without whose outstanding skills in microinjection this work could not have been carried out. The work was funded by a grant from the Simons Initiative for the Developing Brain.

## Supplementary data

**Supplementary figure 1.**
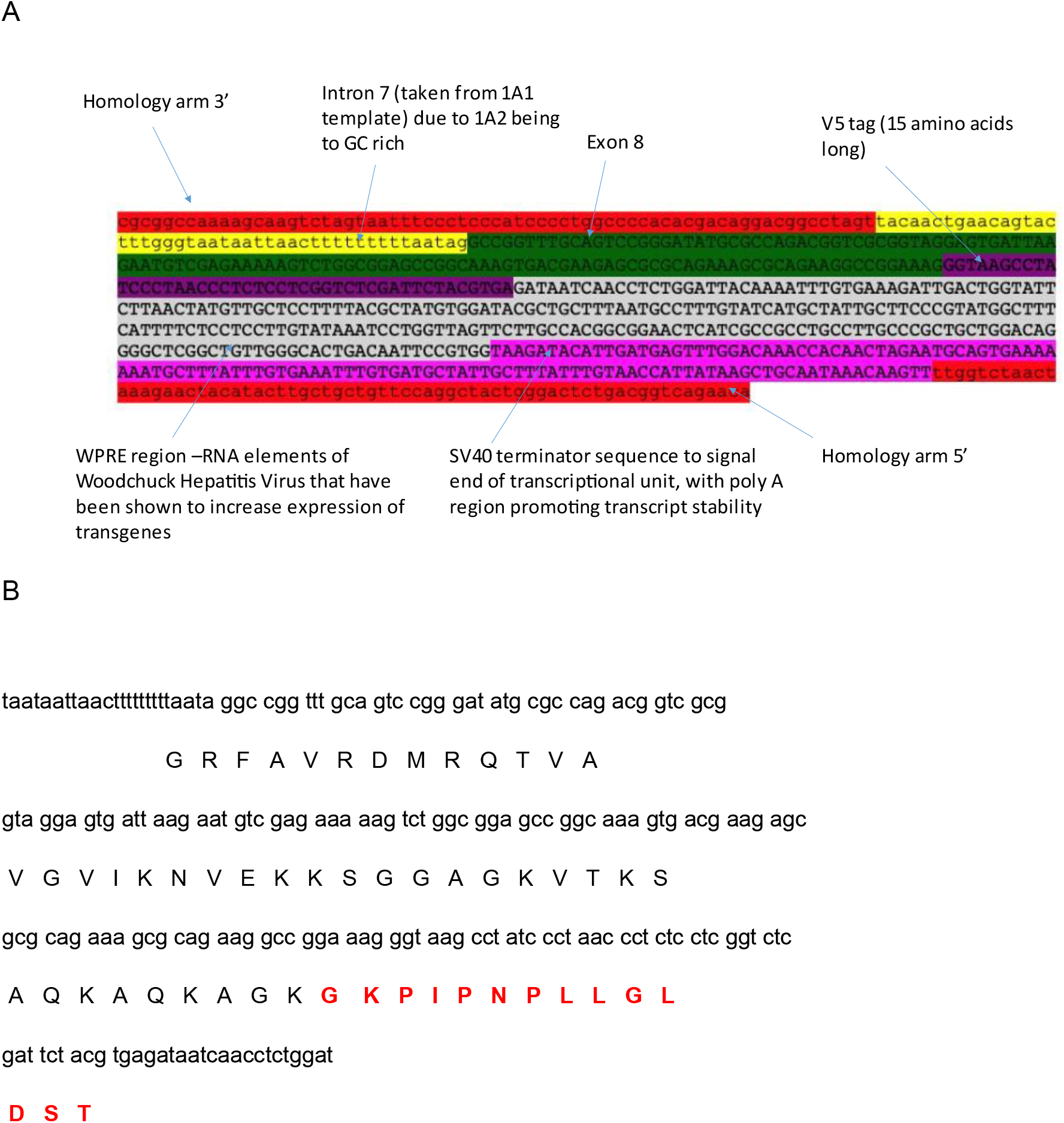
A) Schematic of repair template and B) conceptual translation to show V5 tag (in red) in frame

